# Using topic modeling to detect cellular crosstalk in scRNA-seq

**DOI:** 10.1101/2021.07.26.453767

**Authors:** Alexandrina Pancheva, Helen Wheadon, Simon Rogers, Thomas Otto

## Abstract

Cell-cell interactions are vital for numerous biological processes including development, differentiation, and response to inflammation. Currently most methods for studying interactions on scRNA-seq level are based on curated databases of ligands and receptors. While those methods are useful, they are limited to our current biological knowledge. Recent advances in single cell protocols have allowed for physically interacting cells to be captured and as such we have the potential to study interactions in a complimentary way without relying on prior knowledge. We introduce a new method for detecting genes that change as a result of interaction based on Latent Dirichlet Allocation (LDA). We apply our method to synthetic datasets to demonstrate its ability to detect genes that change in an interacting population compared to a reference population. Next, we apply our approach to two datasets of physically interacting cells and identify genes that change as a result of interaction, examples include adhesion and co-stimulatory molecules which confirm physical interaction between cells. For each dataset we produce a ranking of genes that are changing in subpopulations of the interacting cells. In addition to genes, discussed in the original publications we highlight further candidates for interaction in the top 100 and 300 ranked genes. Lastly, we apply our method to a dataset generated by a standard droplet based protocol, not designed to capture interacting cells and discuss its suitability for analysing interactions. We present a method that streamlines the detection of interactions and does not require prior clustering and generation of synthetic reference profiles to detect changes in expression.

**Author summary:** While scRNA-seq research is a dynamic area, progress is lacking when it comes to developing methods that allow analysis of interaction that is independent of curated resources of known interacting pairs. Recent advances of sequencing protocols have allowed for interacting cells to be captured. We propose a novel method based on LDA that captures changes in gene expression as a result of interaction. Our method does not require prior information in the form of clustering or generation of synthetic reference profiles. We demonstrate the suitability of our approach by applying it to synthetic and real datasets and manage to capture biologically interesting candidates of interaction.

## Introduction

Over the past years single cell RNA-seq (scRNA-seq) has gained immense popularity as it provides the capability to study rare cell types, heterogeneity of cell populations, and disease and developmental trajectories [1].

Cell-cell communication is vital for most biological processes, from maintaining homeostasis to determining specific immune responses [2]. Malfunctioning cells can induce secondary changes in their micro-environment which leads to reprogramming of the niche to their advantage [3]. Improving understanding of essential interactions has the potential to aid discovery of novel therapeutic targets [4].

However, for cell-cell interaction most scRNA-seq studies focus on curated resources of known interacting pairs. Examples of such methods, using a priori curated interactions include: CellPhoneDb, NicheNet, and SingleCellSignalR [5], [6], [7]. CellPhoneDB or variations of their method have been applied in practice to answer questions about inter-cellular communication between cell types in a range of tissues. For example, Cohen et al [8] consider the interaction of lung basophils with the immune and non-immune compartment by examining known ligand-receptor pairs and how those potentially link to development. As these methods are based on databases, they do not allow for new interactions to be identified and results are limited to known biology. Furthermore, most curated resources of ligands-receptor pairs are only available for humans or mouse ortologs [5].

In scRNA-seq, it is possible for two cells to be sequenced together, known as “doublets”. Often doublets are a result of errors in cell sorting or capture but recently two studies have shown that doublets can capture two physically interacting cells (PICs), offering a valuable method to measure, without relying on prior knowledge, the transcription pattern of interaction. Boisset et al [9] used mouse bone marrow (BM) to demonstrate that cell-cell interaction can be studied by dissociating physically interacting doublets. The two interacting cells are separated by needles and sequenced. Further experiments also managed to infer interactions by sequencing intact doublets which were then deconvoluted based on the gene expression of the sequenced singlets. Giladi et al [10] developed a method for sequencing PICs, known as PIC-seq. With other single cell technologies, information about cell-cell interactions are lost due to cell dissociation while PIC-seq captures pairs of interacting cells. PICs are isolated by a combination of tissue dissociation, staining for mutually exclusive markers, and flow cytometry sorting. Single positive and PIC populations are then sequenced. The ability to capture PICs allows Giladi et al [10] to go beyond interactions described in curated databases and potentially identify novel ones. On the computational side of their PIC-seq approach, they [10] cluster the mono-cultures and from these the gene expression of each PIC is modeled as a doublet: *α* \**A* + (1 − *α*) * *B* where *A* and *B* are the two cell types that make the PIC and *α* is the mixing parameter. *α* is estimated by a linear regression model trained on synthetic PICs. This is followed by maximum likelihood estimation (MLE) of *A* and *B*. By identifying what the two subtypes that make the PIC are, expected expression can be computed. Expected and actual expression of the PIC are compared to identify changes as a result of interaction [10]. There are several potential limitations of the outlined approach. Since the PIC-seq algorithm relies on deconvoluting doublets, it cannot be applied to transcriptionally similar cells, such as subtypes or same cell type. Furthermore, for the training of the linear regression, synthetic PICs are created by pairing pooled cells from *A* and *B* that are then downsampled to a predefined total number of unique molecular identifiers (UMIs). While the approach of combining cell profiles to create a doublet has been used, with some modifications, in a range of studies [11], [12], it simplifies how a doublet gets created in practice [13]. Additionally, the method described in PIC-seq requires prior clustering of cells before simulating artificial PICs and deconvolution.

Initially developed for text, LDA has been applied to different types of omics data [14]. In single cell, LDA has been applied to RNA-seq, ATAC-seq, and Hi-C data. In the context of LDA, cells are equivalent to documents and genes (regions in ATAC-seq and locus-pairs in Hi-C) are words. Topics can be described as groups of genes whose expression co-varies [15], [16], [17]. When interpreting the identified topics, they can be labelled as general, cell type specific or linked to technical quality of the samples (e.g ribosomal or mitochondrial dominated topics might correspond to dying cells). In addition to the standard implementation of LDA, work has been completed to allow for simultaneous topic identification and cell clustering [18]. Furthermore, CellTree [19] uses LDA for trajectory inference: the method takes LDA in its standard form but computes chi-square distance between cells and uses the distance to build a tree as a way of describing a branching proces. Most recently, a modified version of LDA has been used to decontaminate counts from ambient RNA: DecontX assumes counts come from two topics, native counts and ambient RNA. Using only the native counts improves clustering and downstream analysis [20].

In this article we propose a novel method based on LDA that allows for identification of genes that change their expression as a result of cell-cell interaction. Once trained on a reference population we can fit LDA on an interacting population and capture changes that cannot be explained by the initially learned topics. Firstly, we show how the proposed model behaves when fit on synthetic doublets with some up-regulated genes. We also show new topics are needed to model the counts of genes related to interaction even if they are not expressed in all interacting cells. We fit LDA as described by [21] on a population of singlets or sorted cells. Then we fix the topics from the singlets reference population and fit another LDA on the interacting cells population. The second LDA allows us to rank the genes that have high probabilities in the new topics. We apply our method to two datasets containing PICs and identify genes that change their expression as a result of interaction, examples of genes include adhesion and co-stimulatory molecules which is a direct evidence of physical interaction between cells. Finally, we demonstrate the challenges of applying our method to a 10x Chromium dataset bronchoalveolar lavage fluid (BALF) of patients with COVID-19 [22]. We link our findings to how well the sequencing protocol can preserve interaction and to what extent we can identify reference populations. However, as the work of [9] and [10] has shown, there is a need to modify currently available scRNA-seq protocols to allow for interactions to be captured. To our knowledge, this is the first paper that models interaction using LDA based approach. Furthermore, our approach does not require prior clustering or synthetic generation of doublets compared to the computational approach previously used to identify genes related to interaction in the work of [10]. We use the genes identified by [10] as a ground truth and show how the number of top genes we select affects true positive and false positive rates. In addition to identifying genes discussed by the original paper, we provide a comprehensive ranking of further genes that might change as a result of interaction such as ones involved in cellular response and adhesion. Taking the top 5 genes per topic in the PIC-seq data allows us to identify 20 further known genes related to cell adhesion and immunity. Additionally, our ranked list of genes includes genes lacking comprehensive annotation and as such allows us to go beyond known interactions. While the analysis of Boisset et al [9] does not consider specific genes that would change as a result of interaction but focuses on cell types known to interact, we perform a literature survey to verify whether we can identify known genes related to interaction in the BM, specifically we consider top 25 genes per topic and we identify over 90 genes linked to cell adhesion and response.

## Materials and methods

### Latent Dirichlet Allocation

Our approach is based on LDA, originally developed by [21] that allows data to be explained by unobserved, latent groups. Those latent groups are called topics and each topic is a multinomial distribution over a set vocabulary. A document can contain words sampled from different topics. This assumption overcomes the limitations of standard mixture models where each document is generated by a single mixture component. As such, LDA allows for documents and biological data to be modeled in a less restrictive way. LDA also allows for different document-topic proportions.

In the context of scRNA-seq, cells correspond to documents and genes to words. Word frequencies are replaced by counts. We obtain a set of topic distributions over cells and a per-topic gene distribution. Given D cells (indexed d = 1,…,D), N genes (indexed n = 1,…,N), and K topics, we can define the generative process as follows:

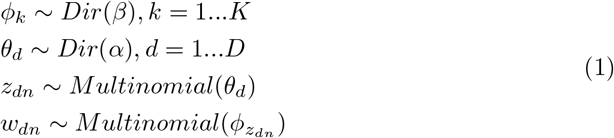

where *α* and *β* are vectors of lengths K and V where V is the size of the vocabulary (all genes in the dataset). *α* and *β* are Dirichlet priors that control the number of topics per cell and number of genes per topic. Those priors control the sparsity of the model.

In our proposed approach, we initially fit LDA on a reference population: co-cultures of a single cell type in the PIC-seq dataset, sorted BM cells in the case of the Boisset’s dataset and what we identify as singlets in the case of the COVID-19 BALF data. The assumptions of LDA fit well in the context of scRNA-seq as at any given point we can observe multiple processes in a cell, as such a cell can be described as a contribution from multiple topics, some specific to a cell type and some shared across all cells. Those processes can be described as genes that co-vary. As words can be in multiple topics, genes can be part of multiple processes. By fitting LDA on the co-cultures of one cell type in a dataset, we obtain for each topic a distribution over genes that we then fix before we fit another LDA on the population of PICs, dissociated BM doublets, or DoubletFinder identified doublets respectively for the three datasets discussed in the results. The initial LDA captures a reference state of cells, a state without interaction. Fixing some of the topics learned from the reference, not interacting populations, allows us to capture in the new topics any changes as a result of interaction due to the setting of the datasets analysed.

We use variational inference to estimate the parameters of the posterior distribution of LDA. As LDA has been extensively discussed in literature, an explanation of the inference procedure can be found in [21].

### Identifying topics linked to a cell type

In order to aid interpretation of the identified topics, we link the topics to celltypes. For each topic we group together cells from the same cell type and perform Mann–Whitney U test. To correct for multiple testing, we use the Benjamini-Hochberg procedure with alpha set to 0.05.

### Finding the number of topics

To select a suitable number of topics, we compute average cosine distance, r, within topics as defined by [23]. A lower value indicates a more stable model. For example, when trying to choose the number of topics for our PIC-seq dataset, we run LDA on the co-cultures several times with the following number of topics: 30, 50, 70, and 100. The table below shows the average cosine distance for each model:

**Table 1.** Average cosine distance between topics for several topic configurations for a model trained on two co-cultures, one of T-cells and another of DCs. Based on the cosine distance, the most suitable number of topics for this dataset would be 30.

| Number of Topics | Average Cosine Distance |
| --- | --- |
| 30 | 0.001842 |
| 50 | 0.002025 |
| 70 | 0.002728 |
| 100 | 0.001874 |

### Motivating the need for new topics

Consider a cell n being one of the PICs. Cell n decomposes into *θ*_*n*_ where Σ_*k*_ *θ*_*nk*_ = 1 (probability distribution for cell n over all k topics). Some of the topics come from the mono-culture LDA fit, those are the topics we fixed before fitting the second LDA, and some topics come from the PICs. We want to compare a fit with all topics with a fit where we do not use any new topics. Let Δ_*nk*_ = *θ*_*nk*_ but we set the contribution of all topic probability for topic k and Σ_*m*_ *β*_*km*_ = 1. If we are only interested in the topics that come from the PICs to 0 and re-normalise, so that Σ_*k*_Δ_*nk*_ = 1. Let *β*_*k*_ = topic probability of picking the counts for one gene versus all other genes, the multinomial distribution reduces to a binomial. For each topic distribution, we compute the probability *P* (*X >*= *x*) for a binomial distribution defined as *X ∼ Binomial*(*n, p*) where n is the total counts for cell n, p is the probability for that gene in that topic, and x is the counts for a particular gene in the current cell. Once we have computed the probability for a gene for each topic, we multiply them by *θ*_*nk*_ and Δ_*nk*_, topic weightings for cell n, and sum on k. We expect the probability of observing counts greater or equal to the actual count for gene changed as a result of interaction to be higher under *θ*_*nk*_ where all topics are included compared to Δ_*nk*_ which is based only on the initial topics. Probabilities should be similar for genes not involved in interaction.

### Ranking genes potential candidates of interaction

One of the outputs from our LDA is a probability distribution for a gene in a cell across all topics. Let N be the doublets/interacting population of interest, then for a gene G and a topic k we find how many cells require topic k to explain the expression of gene G. For a newly identified topic k, for each gene we count how many cells have highest probability for this gene in topic k. For each topic, we produce a ranking of genes based on the number of cells that require this topic to explain the expression of that gene.

The rankings for each newly identified topic can then be analysed. We choose different number of top genes from each topic in subsequent experiments. For example, in the case of our synthetic experiments, we plot how choosing between 2 and 20 genes per topic affect the true positive and false positive rates.

### Evaluation datasets

- PIC-seq of T-cells and DCs: Count matrix and metadata downloaded from the Gene Expression Omnibus (GEO) under accession number GSE135382. Metadata file was used to filter for co-cultures of the same cell type and co-culture of T-cells and DCs. Reference population consists of cells tagged in the metadata as: Co-culture *TCRb*+ (T-cells) and Co-culture *CD*11*c*+ (DCs). All three timepoints for the mono-cultures were used. PICs: Co-culture, *TCRb* + *CD*11*c*+, all three timepoints.
- BM dataset: Count data was acquired from GEO under accession number GSE89379. Sorted cells were used as reference and the dissected doublets were used for analysing interactions. Cells with prefix JC20 to JC47 denote micro-dissected cells. Cells with prefix JC4 denote sorted hematopoietic stem cells (used as reference).
- COVID-19 BALF dataset: Data were downloaded from GEO, accession number GSE145926, in the form of h5 files, CellRanger output. Next subsection describes how reference and potentially interacting populations are identified.

## Pre-processing and analysis before LDA

### PIC-seq and BM dataset

#### PIC-seq

Since the PIC-seq dataset was generated using the MARS-seq platform which has higher sequencing depth than standard 10x Chromium, we set higher filtering cutoffs for number of unique features per cell. As we are not relying on clustering, we can also set a higher cutoff for number of cells a gene is captured in. Genes appearing in fewer than 200 cells were filtered away. Cells with more than 500 features were taken for downstream analysis. Similar to the original publication we exclude ribosomal genes.

#### BM dataset

This dataset has been sequenced using CEL-seq. Only genes with expressed in more than 10 cells were considered for downstream analysis. Resulting in over 10 000 total genes. 369 sorted cells and 1546 dissected doublets were used. No other pre-processing was performed before fitting LDA.

For both datasets filtering steps are performed independently of any scRNA-seq pipeline.

### COVID-19 BALF

Quality control, filtering, normalisation, integration, and clustering were done in R, using Seurat, version 3.1.2. Filtering decisions are dataset dependent and are based on three main metrics: number of genes per cell, number of cells a gene is expressed in, and fraction of mitochondrial genes. It is typical to filter for cells with a very high number of genes expressed to prevent including doublets in the data. For example, cells with low counts and high mitochondrial fraction indicate the mRNA has leaked out through a broken membrane. As such, samples were filtered for cells with fewer than 500 genes. Since we are interested in doublets, which are often assumed to have higher counts than singlets, the maximum cut-off was relaxed [24]. To exclude dying cells we also set a mitochondrial gene expression cut-off to 25. Full list of filtering cut-offs for the different COVID-19 samples can be found in Supplementary Table 1.

Seurat’s NormalizeData and FindVariableFeatures functions were used before integration. Samples were integrated first by condition and then all conditions were integrated using FindIntegrationAnchors. Since compared to the original paper we are not interested in the subtypes of macrophages or T-cells, but the general populations, clustering resolution was set to 0.5. Based on the cluster identification, a subset of the BALF cells were taken forward for the LDA analysis.

To confirm cells we identified as doublets, we use DoubletFinder. Since DoubletFinder can only be used on a single sample and not integrated data, patient samples C143 and C145 were analysed separately [11]. Those two samples were chosen based on amount of cells in what we define as doublet cluster. Supplementary Fig 1 shows the annotated clusters for sample C145. Complete preliminary analysis available at https://github.com/alexpancheva/ldapaper.

## Results and Discussion

### Validation using synthetic doublets

Before testing our method on a real dataset of interacting cells, we apply it on a dataset containing synthetic doublets in order to show that we are able to detect genes that change in interacting cells. We simulate doublets by merging the expression profiles of singlets using different ratios: 50/50 (equal contribution of each cell type), 60/40, and 30/70. In order to obtain a ground truth for genes that change as a result of interaction, we modify the expression of some genes by adding 10 to their total counts. We chose to modify the following genes in the synthetic doublets: (*Sell, Mif, Bcl2l1, Cd40, Myc, Ncl, Cst3, Ly6a, Ctla4, Ccl22, Cd69, Dll4, Lgals1*). We trained our first LDA on a randomly sampled subset of T-cells and DCs mono-culture from the [10] paper. After fixing the topic from the reference, we fit a second LDA on the doublets with upregulated genes. We expect those genes to require contributions from the new topics to model their counts.

We applied the approach described to the datasets of upregulated doublets. We expect the probability of observing counts greater or equal to the actual counts of the list of upregulated genes (e.g. *Ly6a, Sell*) to be higher when we use all topics and we compute the probability under *θ*_*nk*_ where k is topics learned from singlets and simulated interacting doublets. On the other hand, for genes we chose not to upregulate, the probabilities under Δ_*nk*_ and *θ*_*nk*_ should be similar. This is shown in Fig 1, where we plot the probability of observing counts higher or equal to the actual counts in doublets with modified expression of some genes. *Sell*’s counts in the modified doublets can be explained better if the new topics are included. However, in the case of *my-Cytb*, a gene we haven’t modified, probabilities given all topics or only the initial ones are similar.

**Fig 1.**
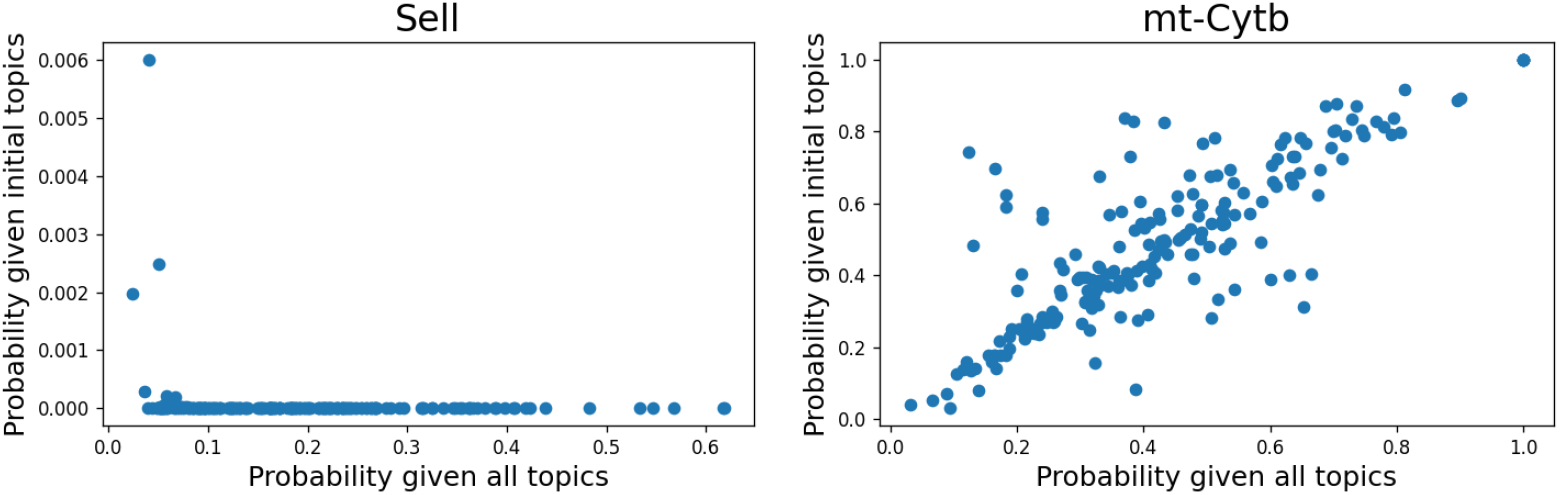
Using the synthetic data, we evaluate how likely it is to observe counts for some genes under *θ* and Δ. For 200 upregulated doublets, we plot the probability of observing the counts of a gene that has been upregulated, *Sell*, and a gene that has not, *mt-Cytb*, under a model using all topics or using only the topics learned from the singlets populations. In the case of *Sell*, a gene with modified expression in the doublets, the probability of observing the actual counts or higher in the upregulated doublets using all topics is higher compared to the probability of observing those counts if we only use the initial topics. However, for *my-Cytb* that we have not upregulated the probabilities under the two models of observing the actual counts or higher are similar. Thus, we can conclude that the additional topics are required to model the genes that change.

For each of the three simulated doublets experiments, we rank the genes based on whether they require contribution from the new topics to explain their expression. We plot true positive rate vs false positive rate using different cutoff values for the top ranked genes. Our final synthetic experiment aims to show we can identify genes that change in a subset of the PICs. We upregulate the expression of *Gbp4, Gbp7, Gzmb, Il2ra, Psma4* in 20 cells (10% of the total PICs). To identify genes of interest we follow the same approach mentioned above and plot a ROC curve, Fig 2, to see how the results are affected by picking a different number of top genes per topic. In all four experiments, using up to 15 top genes per topic resolves the list of upregulated genes we use as ground truth.

**Fig 2.**
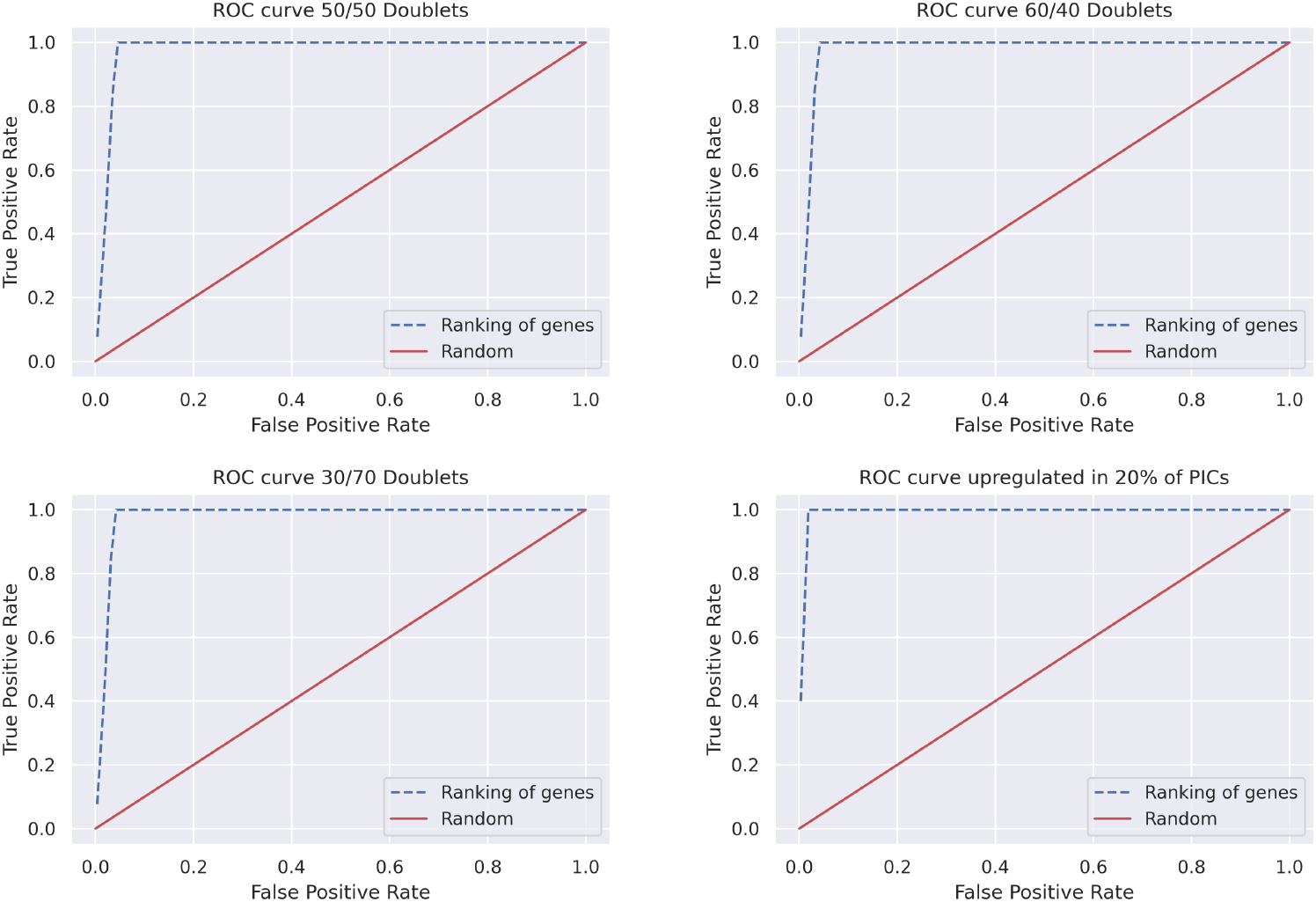
For each of our 4 synthetic experiments, we plot a ROC curve using different number of top genes based on our ranking. In all cases the genes we have modified the expression of appear at the top of the newly identified topics. For all synthetic experiments, using up to top 15 genes per topic resolves the list of upregulated genes we consider as ground truth. However, as it can be seen from the plots even if slightly higher number of top genes used, the false positives are gradually increasing.

### PIC-seq dataset

The first real dataset we use for evaluation has been generated by PIC-seq and includes interacting T-cells and DCs (gated for TCR*β*^+^*CD*11*c*^+^) as well as two co-cultures of a single celltype (T-cells: TCR*β*^+^ and DCs: CD11*c*^+^) across three different timepoints. The original [10] work uses a metacell model to cluster the cultures of a single cell type.

Then each PIC is modeled as a combination of metacells and a mixing proportion, *α*, is estimated by a linear regression model trained on synthetic PICs. The metacells are identified using MLE. Genes of interest are identified by comparing expected expression based on the inferred cell types contributing to the interacting pair and actual expression of the PICs.

Our model does not require prior clustering and generation of synthetic reference profiles and as a first step we train one LDA on the co-cultures of T-cells and DCs using them as a reference. With the first LDA we manage to capture topics specific to T-cells and DCs (groups of genes co-varying in one cell type over the other). As can be seen from Fig. 3 topics 0, 1, 5, and 12 are specific to DCs. We identify topics specific to a cell type as described in Methods. To explore what the top genes are in some of those topics, we pick topics 12 and 23 and order the genes in those topics by probability. We see DC specific genes *Fscn1, Ccl22, Tmem123, and Cd74* in topic 12. Similarly, some of the genes with highest probability in the T-cell specific topic include *Trac, Mif, Ncl, Nmp1* (see Fig. 4).

**Fig 3.**
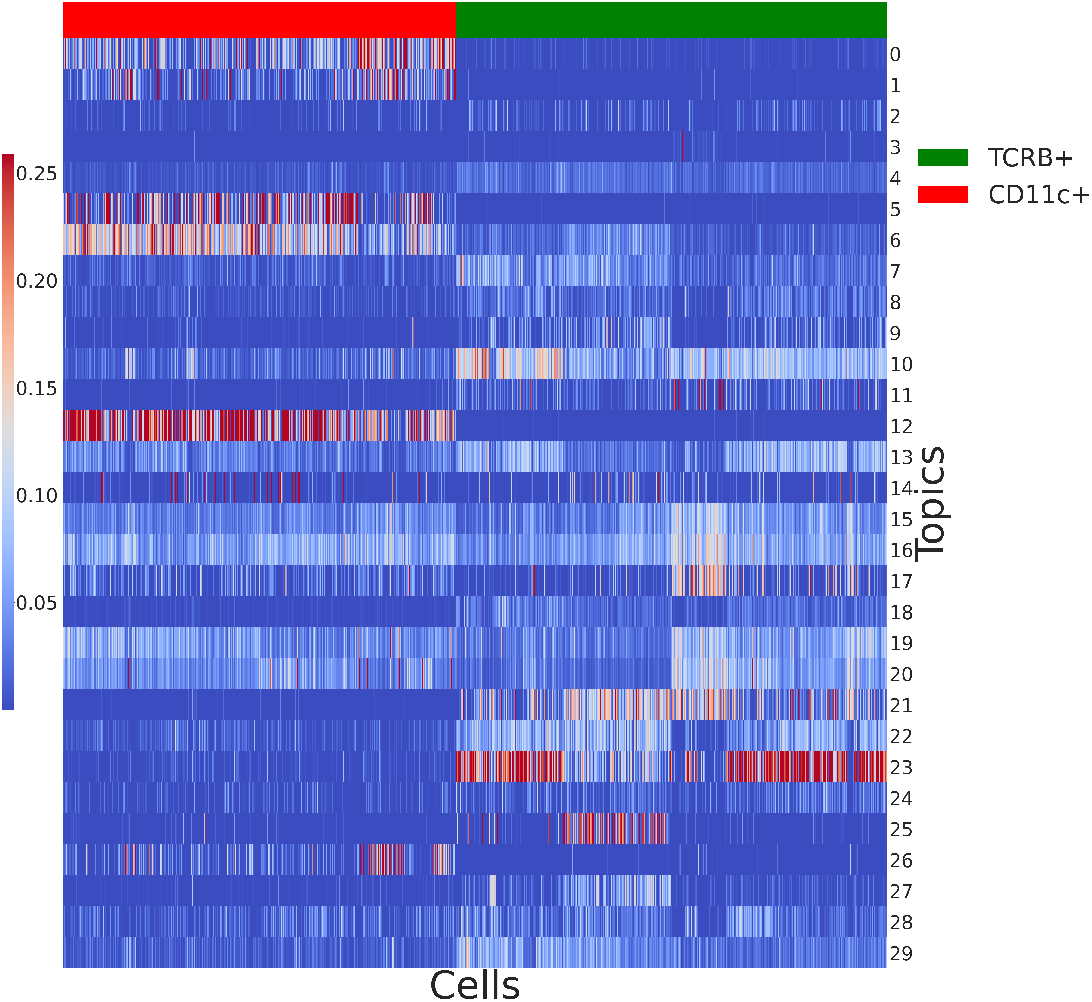
Heatmap of topics expression in the reference populations of T-cells and DCs. To map topics to a cell type, we group together cells from the same cell type and perform Mann–Whitney U test for each topic. Results are corrected using the Benjamini-Hochberg procedure for multiple testing.

**Fig 4.**
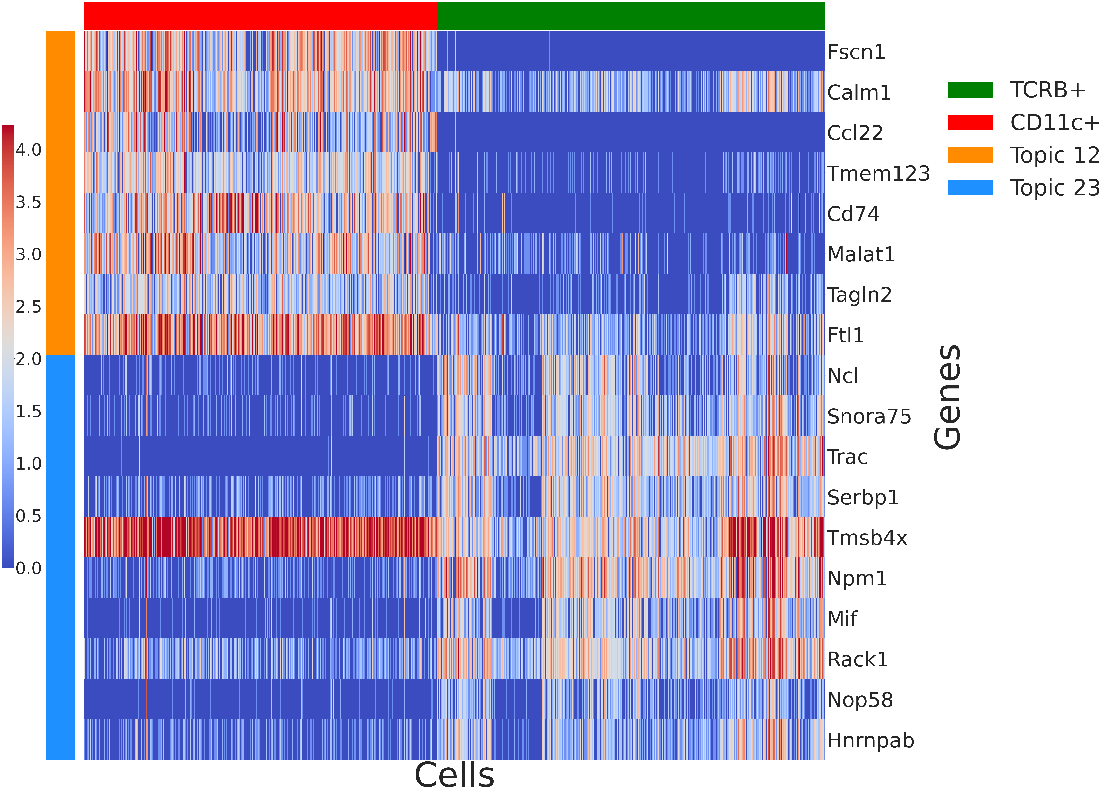
Heatmap of top genes based on probability in that topic (for each topic we can obtain the probability of the genes in that topic) from topics 12 and 23. Topic 12 contains genes that tend to have higher expression in DCs compared to T-cells. Topic 23 contains genes co-varying in the T-cells co-culture. Results similar to Figure 1 from the [10] manuscript

We fix the topics we learned from the co-cultures of the two cell types, T-cell co-culture and DCs co-culture, before we fit another LDA on the physically interacting populations of PICs. As described earlier, in order to rank interesting genes, for each topic we learned from the PICs, we count how many cells use this topic for a particular gene. Then we rank the genes based on number of cells. Our PICs population contains over 3000 cell and we filter our rankings for genes needing particular topic for fewer than 10 cells.

To validate our findings, we check whether the top genes in each of the newly learned topics have also been highlighted by Giladi et al [10] in Supplementary figure 4 of their paper. Due to differences in filtering, we have not retained 10 genes they identify to change as a result of interaction and we take the remaining 81 genes present in our data as a ground truth. In order to evaluate how results are affected by number of top genes per topic we select, we plot true positive rate vs false positive rate, see Fig 5.

**Fig 5.**
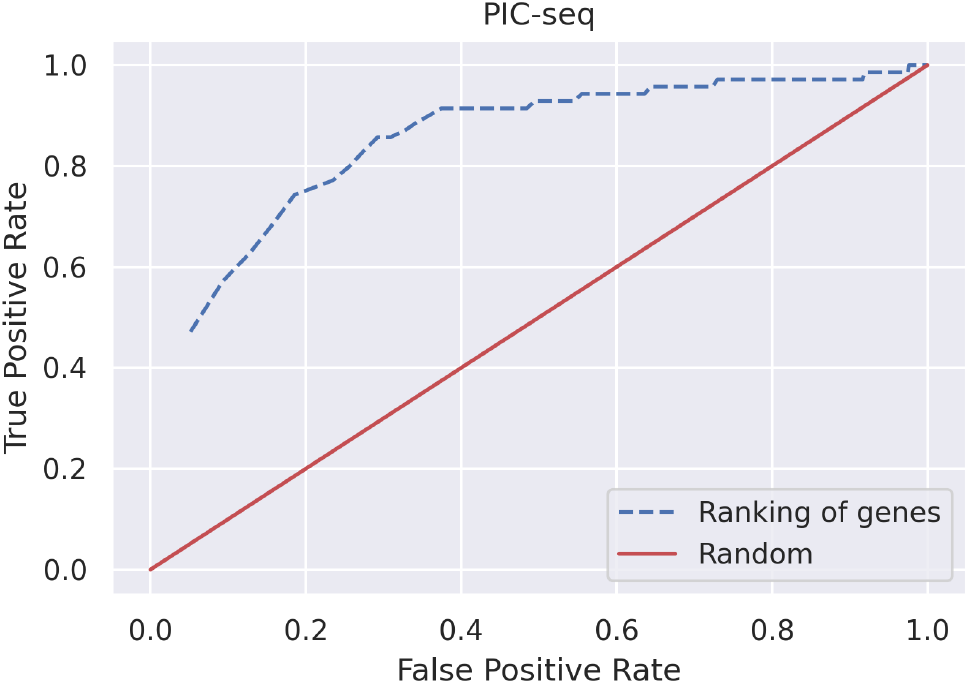
ROC curve using genes from [10] as ground truth based on top genes cutoff for each topic. It’s important to note that some of our false positive values correspond to true interacting genes that have not been presented in the [10] paper amongst their Supplementary fig 4 genes.

While the analysis done in the original publication [10] groups cells by the types of the singlets involved in the interaction and the timepoint of capture before performing log fold change (results in Supplementary figure 4d of the original paper) we show that some of the new topics we identify correspond to the timepoints of capture and reveal genes with temporal patterns as shown on Fig 6. For example, *Ldha, Ptma, Pcna, Trac, Mif, Dut* are needed by a subset of cells and captured in the same topics. Their pattern of expression is higher after the first 3h. *Ncl,Fscn1, Stat4* have higher expression during the first timepoints and appear as top ranked genes for a subset of cells. On the other hand *Tnfrsf4, Tnfrsf9, Tnfrsf18* seem expressed across all timepoints and the shift of their expression is captured by the same topics. The expression of new topics across the cells can be seen on Supplementary Fig 2.

**Fig 6.**
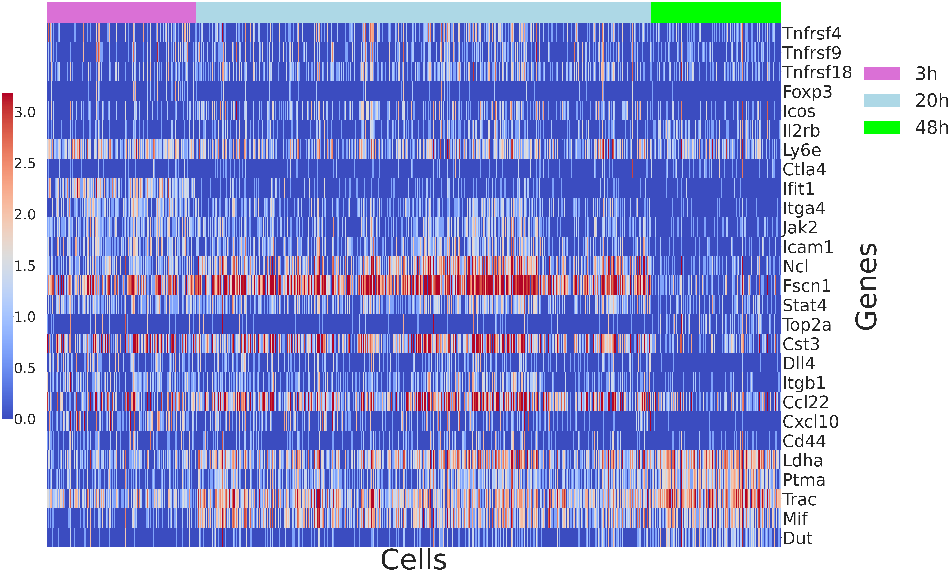
Log transformed expression of genes identified by both [10] and our ranking approach (top 40) and showing the possible temporal expression pattern. For example, genes *Ldha, Ptma, Pcna, Trac, Mif, Dut* have highest probability in the same new topics and their expression increases after the first 3h, while *Tnfrsf4, Tnfrsf9, Tnfrsf18* do not seem to show temporal pattern and the shift in their expression compared to the single cell type co-cultures is captured by the same topics.

While for the purposes of the ROC analysis, we are considering genes not amongst the ones discussed by Giladi et al [10] as false positives, for some of those genes there is evidence they could be involved in interaction. Taking the top 5 genes per topic considered as false negatives previously, we find genes related to immunity and cell adhesion, ligand-receptor pairs (over 20 genes in the first 100 ranked). Examples include *Ighm, Cd48, Ighm, Ly6e*. Additionally, while some known genes appear high in the ranking, some of the genes in our list are not well annotated. This makes them potential targets for further analysis to elucidate their role. Genes can be found in Supplementary Table 2.

### BM dataset

The original work by Boisset et al [9] is focused on identifying significant interactions between cell types, using sorted BM cells and needle dissected doublets. We hypothesised our approach should be able to identify genes involved in the main interactions discussed by [9]. Firstly, we fit our LDA on the sorted BM cells and fix the learned topics. Next, we fit the second LDA on the needle dissected doublets.

We hypothesised our approach should be able to identify genes involved in the main interactions discussed by [9]. Their work looked at three specific interacting pairs: macrophages and erythroblasts, plasma cells and myeloblasts/promyelocytes, and megakaryocytes and neutrophils. Macrophages and erythroblasts have been known to interact and erythroblastic islands are considered an important niche for the maturation of red blood cells. In addition to anchoring erythroblasts within island niches, macrophages also provide interactions important for erythroid proliferation and differentiation [25]. When describing physical interactions, adhesion molecules are of particular interest. In our analysis, we identify *Vcam-1* and *Itgam* which are known to support adhesive interactions in macrophages.

Boisset et al [9] also identify and validate the interaction between megakaryocytes and neutrophils. Their findings support other studies that have looked at emperipolesis (neutrophils are engulfed by BM megakaryocytes) as a process mediated by both lineages. This interaction is important for production of platelets. [26] identified that emperipolesis is mediated by *β*2 integrin Cd18 and Icam-1 interaction. Blocking *β*2 integrin *Cd18 (Itgb2)* impairs emperipolesis [26]. This is another integrin we identify in our analysis. *Elane* and *Igj* are two other genes discussed by Boisset et al [9] that we identify to require additional topics to model their expression. The genes shown in Fig 7 are identified by taking the top 25 genes from each topic. Overall, based on top 25 genes ranking per new topic (over 300 genes in total), we identify genes linked to cell adhesion, innate immunity, and immune response. The full list of genes can be found in the supplementary information, Supplementary Table 2. While some of the genes are known to be linked to neutrophils (*Cd177, Prtn3, Serpinb1a, Lsp1*), other genes are less well-annotated in terms of function and as such this demonstrates the benefits of using an approach that is not based on curated resources of known interactions.

**Fig 7.**
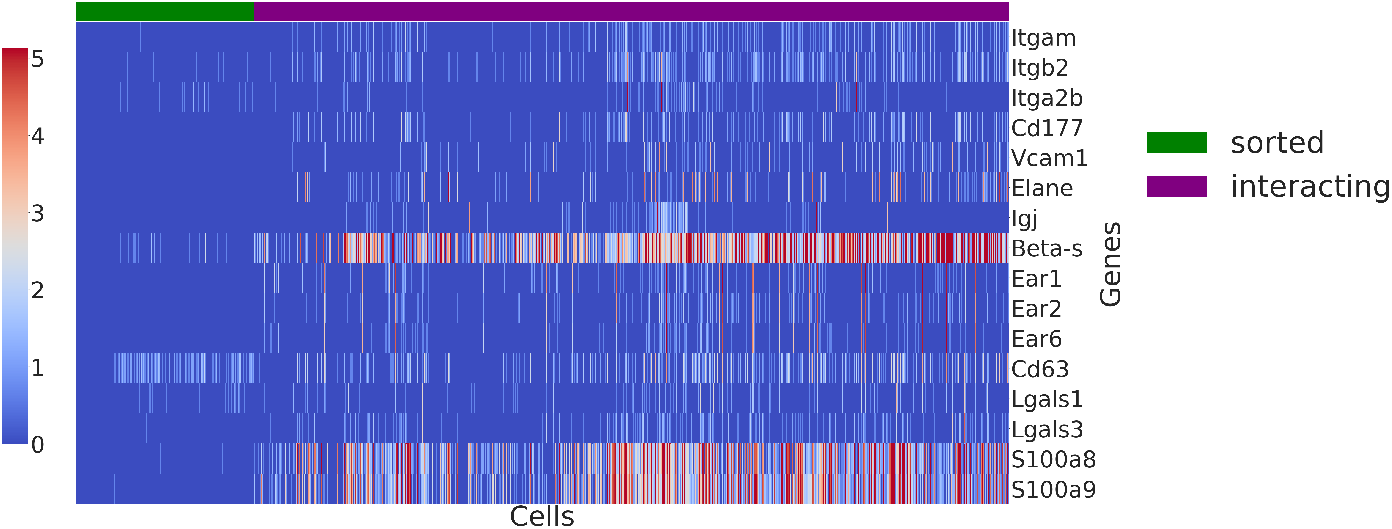
Heatmap of genes chosen based on their ranking in the new topics and evidence from literature that they are linked to interactions in the BM. Amongst the examples we see genes linked to neutrophils adhesion such as *S100a8, S100a9*, and *Cd177. Elane* and *Igj* are two genes confirmed by [9].

### COVID-19 Data

Previously, we used datasets generated by modifying standard protocols to allow for PICs to be generated. However, here we explore the ability of our method to identify genes that change as a result of interaction in datasets generated with the 10x Chromium platform which does not have the ability to preserve interacting cells as there is no specific way of capturing doublets. We took a recently published COVID-19 BALF dataset containing several cell types like T-cells, macrophages, B-cells, DCs, and neutrophills. There are samples from patients with moderate COVID-19, severe infection, and healthy controls. During cluster annotation, the authors labelled several clusters as doublets [22]. We hypothesised some of those doublets might represent interaction as macrophages are known to interact with T-cells. To confirm the identity of the doublet cluster in the severe patient samples, we looked at marker gene expression followed by analysis with DoubletFinder.

We use the populations labelled as singlets by DoubletFinder as reference for LDA. We fix those topics and fit a second LDA on the doublets population. Identification of potential interacting genes was performed similarly to the datasets analysed earlier by ranking genes within each new topic based on how many cells require this particular topic to explain the gene expression. Additionally, as we only have just over 200 doublets, we only consider genes using a specific topic in at least 10 cells. As we can see from Fig 8, some of the genes that require contribution from the new topics to model their expression include cytokines and chemokines which might suggest interaction. A subset of the cells also seem to require new topics for genes related to interaction. We refer to work by Takada et al [27] and Magee et al [28] that discuss physical interactions to identify suitable candidates. As not all doublets require new topics, it is possible we have a mix of interacting and technical doublets. While we are capturing a shift in the expression of some genes, our results are inconclusive due to the quality of our reference population as the reference is constructed based on the cells DoubletFinder annotated as singlets. Depending on the amount of cells loaded, a standard 10x Chromium experiment can result to the formation of over 0.7% technical multiplets with doublets being predominant. While with DoubletFinder we have managed to label some of the doublets, computational tools for doublet detection do not achieve perfect sensitivity and specificity scores, so it is possible the reference population contains technical doublets or interaction cells. Our approach can identify genes that change as a result of interaction and is suitable for datasets where the reference population is clearly labelled as in the PIC-seq and dissociated BM examples discussed earlier.

**Fig 8.**
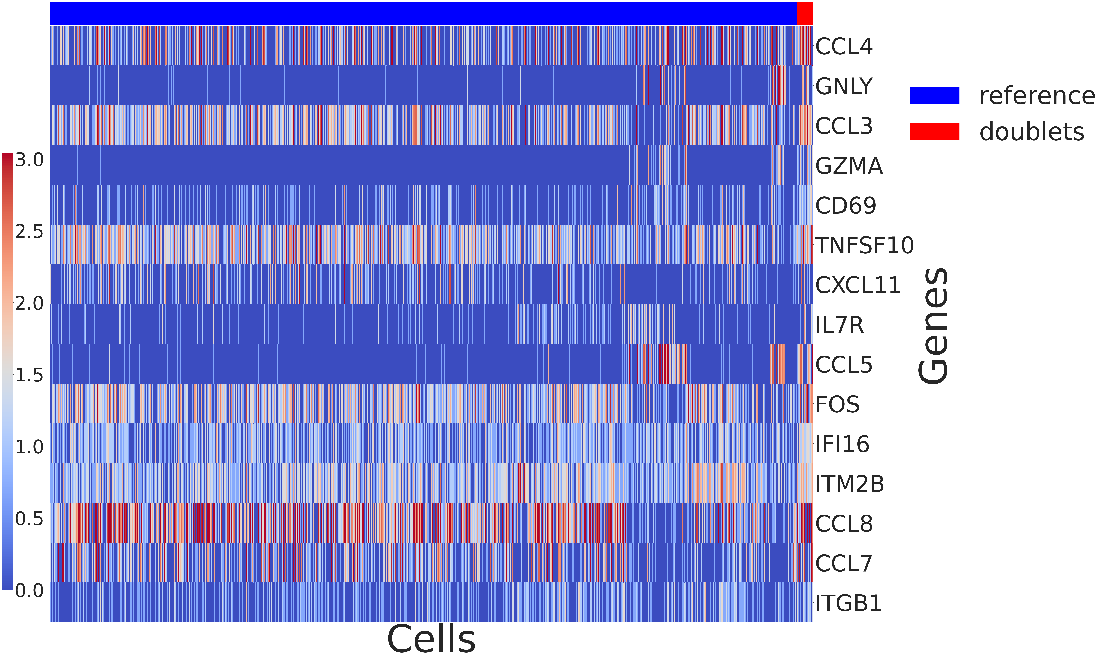
Heatmap of the COVID-19 data showing the expression of top ranked genes needing new topics for a subset of the doublets population. As none of genes are uniquely expressed in the doublets population, we are capturing a shift in the expression of those genes. While some doublets can represent biologically interacting ones as top ranking genes include cytokines and chemokines, due to the quality of the reference population our results are inconclusive.

## Conclusion

In this article we have demonstrated the suitability of LDA for analysing PICs. We have shown that our model is sensitive to changes in gene expression that cannot be explained by the not interactive populations and thus new topics are needed to model the expression of genes that change as a result of interaction. Our model has been applied to two datasets of sequenced PICs and a dataset generated by standard 10x Chromium. Our approach assumes there is a reference population that can be used to fit the first LDA; for example this could be populations before an interaction has occurred. In addition to genes known to be involved in interaction and discussed by [9] and [10], we also rank further candidates for interaction that might be of interest for validation. We demonstrate the challenges of applying our approach to a dataset where a reference population cannot be clearly labelled in the case of the COVID-19 BALF analysis as with the standard 10x Chromium protocol. However, amongst the top genes we rank there are cytokines and chemokines, genes known to be regulated by physically interacting cells, which might suggest the doublet population includes both technical and interacting doublets. While this is informative, the current setup of the 10x Chromium protocol is fully suitable for studying cellular crosstalk of physically interacting cells and as such the likes of PIC-seq should be considered when studying interactions.

As seen from the datasets discussed here, modeling interactions based on doublets can be very useful. As such, distinguishing technical from biologically significant doublets poses an interesting challenge. While we have applied our LDA approach to PICs, there is a potential for our work to allow for distinguishing technical doublets from transitioning cells as long as the transition is described by a unique set of genes, so that a new topic can be defined. While there are cell hashing methods that allow for mitigation of batch effects and overloading of sequencers, those methods help identify doublets between different samples/patients while intra-donor doublets remain undiscovered. On the computational side of doublet detection, methods make a range of assumptions that pose challenges to using them in practice. For example, DoubletFinder requires doublet rate which is not available in all experiments. The accuracy of all methods is affected when applied to transcriptionally similar cells and DoubletDecon would not allow a doublet cluster to be present in the data. DoubletDecon is the only one able to distinguish technical doublets from transitioning cells. As such, there is a clear need for methods that can eliminate technical noise but not at the expense of biological significance [11], [12], [29], [30], [31].

As PIC-seq is a very powerful approach, it can be used to potentially generate data including physically interacting doublets as well as singlets. It would be of interest to identify whether some of the singlets in fact show signs of interaction. Are they cells that have interacted but separated? Or maybe they have not interacted at all? Such datasets could easily be analysed following the methodology described earlier. We would expect to see signs of interacting topics in some singlets but maybe not all and as such we should be able to distinguish between singlets and singlets that have interacted before.

As there is a demand for understanding PICs, we believe methods like PIC-seq will be used more often in future and further work will be done to develop sequencing protocols that allow for capturing physical interactions that have the potential to become therapeutic targets. When such datasets are generated, there should be techniques that allow for their analysis and are not limited to knowledge captured in biological databases. The method described here is one such example that does not require any prior information such as clustering of the cell types involved and generation of synthetic reference profiles. As further such datasets are generated, fields that would benefit from more in depth understanding of interactions include: understanding parasite-host interactions, crosstalk between immune cells and other lineages, and effect of cell-cell interaction in cancer progression.

## Supporting information

**S1 Table.** Filtering parameters used for each sample in the COVID-19 dataset. Upper cutoffs for nFeatures has been set to relatively high values as we are interested in potential doublets. The percent.mt (% mt) cutoff allows us to exclude dying cells.

| Sample Name | features lower cutoff | features upper cutoff | % mt |
| --- | --- | --- | --- |
| C51 | 500 | 3000 | 25 |
| C52 | 500 | 2000 | 20 |
| C100 | 500 | 6000 | 25 |
| C141 | 500 | 7000 | 25 |
| C142 | 500 | 6500 | 25 |
| C144 | 500 | 4000 | 25 |
| C143 | 500 | 6000 | 25 |
| C145 | 500 | 4500 | 20 |
| C146 | 500 | 4500 | 25 |
| C148 | 500 | 7500 | 25 |
| C149 | 500 | 7000 | 25 |
| C152 | 500 | 6000 | 25 |

**S1 Fig.**
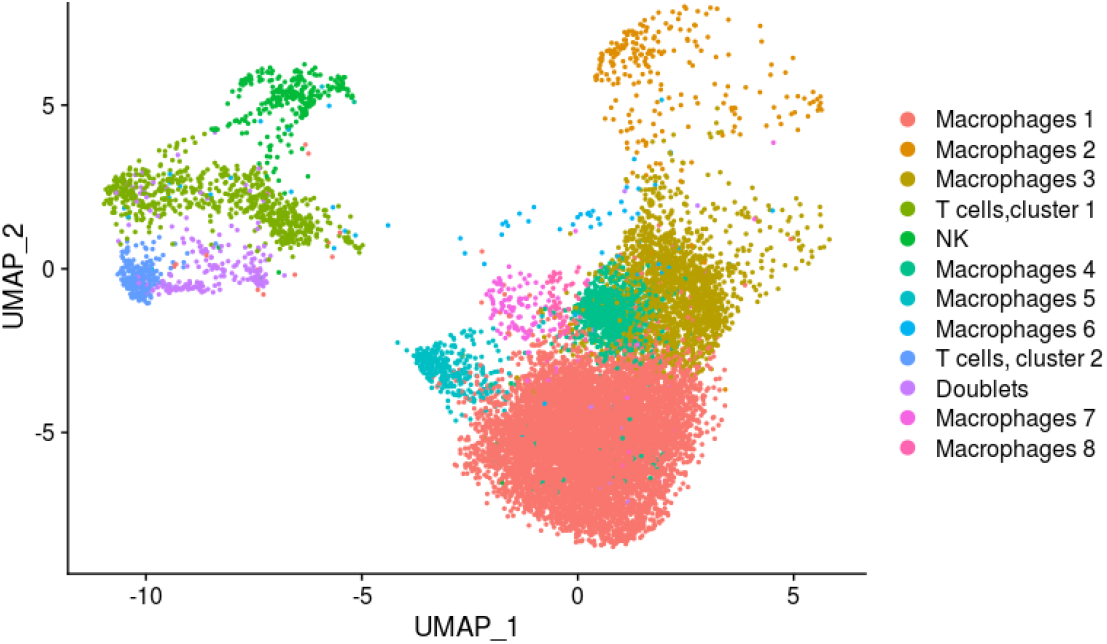
C145 cluster annotation: COVID-19 BALF, UMAP projection of patient sample C145. We have identified the cluster containing doublets based on expression of marker genes and annotation by DoubletFinder.

**S2 Table Added as a separate table: Top genes from the PIC-seq and the bone marrow datasets**

**S2 Fig.**
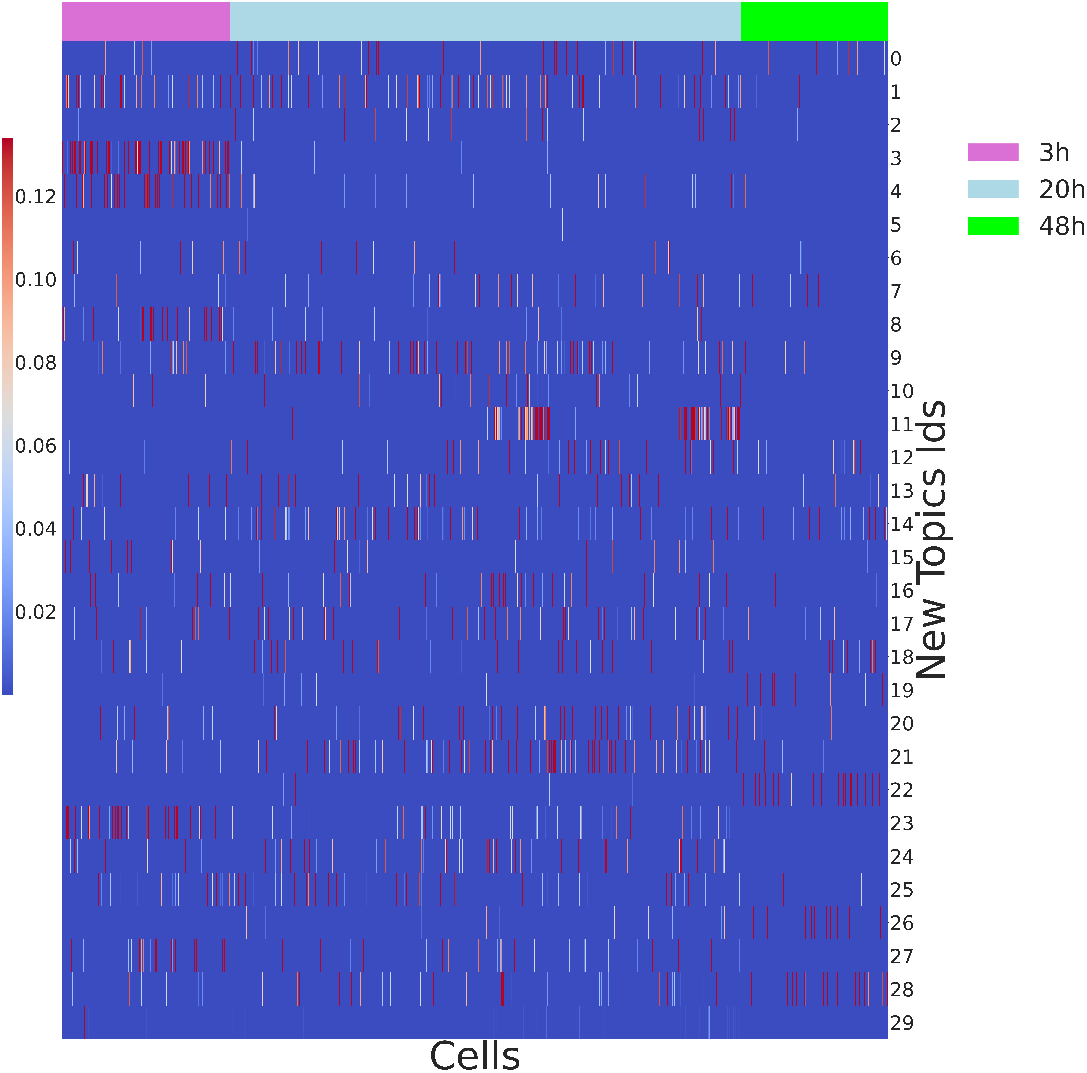
New topics and PICs heatmap. While some topics have higher probabilities associated with a timepoint. Other topics appear expressed across all timepoints, so the gene expression shift is linked to the cell types that contribute to the PICs.

## Acknowledgments

**AP is supported by MRC grant number: MR/N013166/1. TO is supported by the Wellcome Trust grant 104111/Z/14/ZR**.

